# Interdependent Allosteric FFA2R Modulators Synergistically Induce Functional Selective Activation and Desensitization in Neutrophils

**DOI:** 10.1101/809061

**Authors:** Simon Lind, André Holdfeldt, Jonas Mårtensson, Martina Sundqvist, Terry P. Kenakin, Lena Björkman, Huamei Forsman, Claes Dahlgren

## Abstract

The non-activating allosteric modulator AZ1729, specific for free fatty acid receptor 2 (FFA2R), transfers the orthosteric FFA2R agonists propionate and the P2Y_2_R specific agonist ATP into activating ligands that trigger an assembly of the neutrophil superoxide generating NADPH-oxidase. The homologous priming effect on the propionate response and the heterologous receptor cross-talk sensitized ATP response mediated by AZ1729 are functional characteristics shared with Cmp58, another non-activating allosteric FFA2R modulator. In addition, AZ1729 also turned Cmp58 into a potent activator of the superoxide generating neutrophil NADPH-oxidase, and in agreement with the allosteric modulation concept, the effect was reciprocal in that Cmp58 turned AZ1729 into a potent activating allosteric agonist. The activation signals down-stream of FFA2R when stimulated by the two interdependent allosteric modulators were biased in that, unlike for orthosteric agonists, the two complementary modulators together triggered an activation of the NADPH-oxidase, but not any transient rise in the cytosolic concentration of free Ca^2+^. In addition, following AZ1729/Cmp58 activation, signaling by the desensitized FFA2Rs was functionally selective in that the orthosteric agonist propionate could still induce a transient rise in intracellular Ca^2+^. The novel neutrophil activation and receptor down-stream signaling pattern mediated by the two cross-sensitizing allosteric FFA2R modulators represents a new regulatory mechanism that controls receptor signaling.

**Significance Statement:** A novel activation mechanism of a G-protein coupled free fatty acid receptor (FFA2R), is synergistically triggered by two otherwise non-activating allosteric modulators in the absence of orthosteric agonists. The receptor down-stream signaling proved to be functionally selective (biased); a superoxide generating enzyme is assembled and activated without involvement of the Ca^2+^ signaling pathway. The novel activation mechanism and the receptor down-stream signaling pattern mediated by the two cross-sensitizing allosteric FFA2R modulators, represents a new regulatory mechanism for control of GPCR-signaling.

## Introduction

Plasma membrane spanning G-protein coupled receptors (GPCRs) expressed in many different cell types, typically change their conformation when activating agonists bind to the so called orthosteric binding site, present on the surface of the receptor-expressing cells (Erlandson et al., 2018; Laschet et al., 2018). The conformational change affects the cytosolic domains of the occupied receptor, and initiates an activation of down-stream signaling pathways on the cytosolic side of the receptor expressing membrane. Signaling by GPCRs was long thought to be either on or off depending on whether an agonist was bound or not but, recent research show that the receptor signaling states are more variable as is the classification of the ligands that interact and regulate receptor functions (Smith et al., 2018; Wang et al., 2018); conventional agonists activate the targeted receptor and amplify multiple classical signaling GPCR cascades, whereas other ligands give rise to functionally selective responses (Smith et al., 2018). The molecular basis for such a biased/functional selective response is illustrative for the structural diversity and variability of agonist occupied receptors, allowing receptors to be stabilized in a conformation that promotes either a balanced response or one signaling pathway over another (Smith et al., 2018). In addition, receptor selective ligands may have alternative (allosteric) binding sites distinctly separated from the orthosteric site both physically and structurally. Ligands, that bind to such regulatory receptor sites, are commonly neutral without a concomitant binding of a conventional (orthosteric) agonist, yet modulate the signaling properties of targeted receptors when activated by a conventional agonist (Gentry et al., 2015). This mechanism, referred to as allosteric receptor modulation, by which a receptor is transferred to a sensitized state by one ligand when activated by another conventional orthosteric agonist, should, according to the allosteric GPCR-modulation dogma, affect signaling properties solely of ligands that interact specifically with the allosterically modulated receptor (Changeux and Christopoulos, 2016; Kenakin, 2017a). This is, however, not always the case; recently, an allosteric modulator (Cmp58) specific for the short chain free fatty acid 2 (FFA2R; (Wang et al., 2010)), was shown to affect signaling mediated not only by orthosteric agonists but also by an agonist for the ATP-receptor P2Y_2_R and for the formyl peptide receptor 2 (FPR2; (Lind, 2019a). The recent data presented demonstrate that natural FFA2R agonists trigger an increase in the cytosolic concentration of free Ca^2+^ in neutrophil phagocytes, an activation achieved without any assembly of an enzyme system (the phagocyte NADPH-oxidase) designed to generate superoxide anions (Lind, 2019a; Martensson et al., 2018). In accordance with the allosteric receptor modulation concept, the allosteric FFA2R modulator Cmp58 lacked direct activating effect on neutrophils but acted as a positive modulator that turned FFA2R agonists into potent NADPH-oxidase activating ligands. In addition, the responses induced by ATP and fMLF were also primed by the modulator (Lind, 2019a; Martensson et al., 2018). It is, thus, obvious that the down-stream signaling and non-specific receptor effects of allosteric receptor modulators are much more complex than initially anticipated.

A novel allosteric FFA2R ligand, AZ1729, was recently described and shown to be both a positive allosteric FFA2R modulator that increased the activity induced by conventional orthosteric agonists and a direct activating allosteric agonist (Bolognini et al., 2016). It is also interesting to note, that the allosterically modulated FFA2Rs were able to signal through different G-proteins containing the G*α*i and G*α*q subunits, respectively (“induced-bias” (Bolognini et al., 2016; Kenakin, 2015).

It is reasonable to assume that FFA2R has multiple ligand binding sites allowing different allosteric modulators to distinctly affect the affinity/efficacy of orthosteric ligands. We have now investigated the effects of AZ1729 on neutrophil FFA2R-signaling and show that this novel allosteric FFA2R modulator affects neutrophil function/signaling in the same way as the earlier described allosteric FFA2R modulator Cmp58. That is, it lacks direct activating effects in its own but turns the natural FFA2R agonist propionate into a potent activator of the neutrophil NADPH-oxidase, and has off-target receptor cross-talk effects on the receptors for ATP (P2Y_2_R) and formylated peptides (FPRs; (Lind, 2019b)). More importantly, we show that binding of the non-activating allosteric modulator AZ1729 sensitized FFA2R also to the other non-activating allosteric modulator Cmp58. The novel activation/sensitization mediated by the two allosteric FFA2R modulators was reciprocal, and represents a new regulatory mechanism that controls GPCR signaling. In addition, the down-stream signaling of FFA2R was biased in that the two interdependent modulators activated the neutrophil NADPH-oxidase, but no transient rise in the free cytosolic concentration of calcium ions (Ca^2+^) was induced. Signaling by the AZ1729/Cmp58-desensitized FFA2Rs also displayed functional selectivity in that the orthosteric FFA2R agonist propionate retained the ability to induce a transient rise in intracellular Ca^2+^.

## Material and Methods

### Chemicals

Isoluminol, TNF-*α*, ATP, propionic acid, bovine serum albumin (BSA), and fMLF were purchased from Sigma (Sigma Chemical Co., St. Louis, MO, USA). Cyclosporin H was a kind gift provided by Novartis Pharma (Basel, Switzerland). Dextran and Ficoll-Paque were obtained from GE-Healthcare Bio-Science (Uppsala, Sweden). Fura 2-AM was from Molecular Probes/Thermo Fisher Scientific (Waltham, MA, USA), and horseradish peroxidase (HRP) was obtained from Boehringer Mannheim (Mannheim, Germany). The allosteric FFA2R modulator AZ1729 (Bolognini et al., 2016) was a generous gift from AstraZeneca (Mölndal, Sweden) and the phenylacetamide compound (S)-2-(4-chlorophenyl)-3,3-dimethyl-N-(5-phenylthiazol-2-yl)butanamide (PA;Cmp58 (Wang et al., 2010)) was obtained from Calbiochem-Merck Millipore (Billerica, USA) The Gαq inhibitor YM-254890 was purchased from Wako Chemicals (Neuss, Germany). The FFA2R antagonist CATPB ((S)-3-(2-(3-chlorophenyl)acetamido)-4-(4-(trifluoromethyl)-phenyl) butanoic acid) synthesized as described previously (Due-Hansen et al., 2015; Hudson et al., 2013; Hudson et al., 2012), was a generous gift from Trond Ulven (Odense university, Denmark). The P2Y_2_R antagonist AR-C118925XX (5-[[5-(2,8-Dimethyl-5*H*-dibenzo[*a*,*d*]cyclohepten-5-yl)-3,4-dihydro-2-oxo-4-thioxo-1(2*H*)-pyrimidinyl]methyl]-*N*-2*H*-tetrazol-5-yl-2-furancarboxamide) was obtained from Tocris (Abingdon, UK). Subsequent dilutions of receptor ligand and other reagents were made in Krebs-Ringer Glucose phosphate buffer (KRG; 120 mM NaCl, 4.9 mM KCl, 1.7 mM KH_2_PO_4_, 8.3 mM NaH_2_PO_4_, 1.2 mM MgSO_4_, 10 mM glucose, and 1 mM CaCl_2_ in dH_2_O, pH 7.3).

### Isolation of human neutrophils

Neutrophils were isolated from buffy coats from healthy blood donors by dextran sedimentation and Ficoll-Paque gradient centrifugation as described by Bøyum (Boyum et al., 1991). Remaining erythrocytes were removed by hypotonic lysis and the neutrophils were washed and resuspended in KRG. To amplify the activation signals the neutrophils were primed with TNF-*α* (10nM for 20 min at 37°C), and then stored on ice until use.

### Measuring NADPH-oxidase activity

Isoluminol-enhanced chemiluminescence (CL) technique was used to measure superoxide production, the precursor of production of reactive oxygen species (ROS), by the NADPH-oxidase activity as described (Bylund et al., 2014; Dahlgren and Karlsson, 1999). In short, the measurements were performed in a six-channel Biolumat LB 9505 (Berthold Co., Wildbad, Germany), using disposable 4-ml polypropylene tubes and a 900 µl reaction mixture containing 10^5^ neutrophils, isoluminol (2×10^−5^ M) and HRP (4 Units/ml). The tubes were equilibrated for 5 min at 37°C, before addition of agonist (100 µl) and the light emission was recorded continuously over time. In experiments where the effects of receptor specific antagonists were determined, these were added to the reaction mixture 1-5 min before stimulation with control neutrophils incubated under the same condition but in the absence of antagonist run in parallel for comparison.

### Calcium mobilization

Neutrophils at a density of 1–3×10^6^ cells/ml were washed with Ca^2+^-free KRG and centrifuged at 220x*g*. The cell pellets were re-suspended at a density of 2×10^7^ cells/ml in KRG containing 0.1% BSA, and loaded with 2 μM FURA 2-AM for 30 min at room temperature. The cells were then washed once with KRG and resuspended in the same buffer at a density of 2×10^7^cells/ml. Calcium measurements were carried out in a Perkin Elmer fluorescence spectrophotometer (LC50), with excitation wavelengths of 340 nm and 380 nm, an emission wavelength of 509 nm, and slit widths of 5 nm and 10 nm, respectively. The transient rise in intracellular calcium is presented as the ratio of fluorescence intensities (340 nm / 380 nm) detected.

### The functional allosteric model used to describe the FFA2R allosterism

Allosteric parameters describing the co-operativity between AZ1729 and Cmp58 were obtained by fitting dose-response data to a functional allosteric model as described (Martensson et al., 2018). In short, the model combines the Stockton-Ehlert allosteric binding model (Ehlert, 1988; Stockton et al., 1983) with the Black/Leff operational model (Black and Leff, 1983) for functional activity to describe allosterism; the effects of an allosteric modulator on agonist function (Ehlert, 2005; Kenakin, 2005; Price et al., 2005). The model allows the receptor R to bind the two ligands [A] and [B] concomitantly; function is produced through the [AR] and/or [ABR] complexes interacting with cellular components according to the Black/Leff operational model to yield cellular response. The model yields an equation to predict the response to the agonist in the presence of the modulator (Ehlert, 2005; Kenakin, 2005; Price et al., 2005):

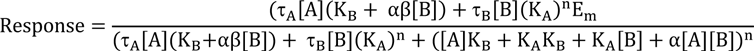

This analysis yields values that characterize the allosteric interaction of [A] and [B] in a system independent manner. The allosteric effect is given by discrete values of *α* (effect on affinity), *β* (effect on efficacy), K_B_ (equilibrium dissociation constant of the B in the absence of A) and where applicable, *τ*_B_ (the efficacy of the modulator). What is required is the efficacy of the agonist (*τ*_A_), n (slope of the concentration-response curves) and some measure of the maximal response capability of the system (E_m_); these are obtained through fitting the concentration response curve to [A] in the absence of [B] as done routinely when quantifying the response to an agonist through the Black/Leff operational model (Black and Leff, 1983; Black et al., 1985).

### Statistical analysis

Statistical calculations were performed in GraphPad Prism 8.02 (Graphpad Software, San Diego, CA, USA). The specific statistical tests are stated in the relevant figure legend. A p-value < 0.05 was regarded as statistically significant difference and is indicated by *p < 0.05, **p < 0.01. Statistical analysis was performed on raw data values using a one-way ANOVA followed by Dunnett’s multiple comparison or, paired Student’s t-test.

### Ethics Statement

In this study, conducted at the Sahlgrenska Academy in Sweden, we used buffy coats obtained from the blood bank at Sahlgrenska University Hospital, Gothenburg, Sweden. According to the Swedish legislation section code 4§ 3p SFS 2003:460 (Lag om etikprövning av forskning som avser människor), no ethical approval was needed since the buffy coats were provided anonymously and could not be traced back to a specific donor.

## Results

### Two allosteric FFA2R modulators, AZ1729 and Cmp58, have similar sensitizing effects on neutrophils

#### Allosteric modulation of the propionate induced rise in the concentration of intracellular Ca^2+^[Ca^2+^]_i_

No direct neutrophil activating effect was induced by AZ1729 (structure shown in Fig 1A), as shown by a lack of a change in [Ca^2+^]_i_ (Fig 1C). This is in agreement with the initial description of AZ1729 as a positive specific allosteric FFA2R modulator (Bolognini et al., 2016), as well as with the results obtained earlier with another allosteric FFA2R modulator, Cmp58 (Fig 1B and D; (Lind, 2019a; Martensson et al., 2018)). Further, AZ1729 positively modulated the response induced by propionate, an orthosteric FFA2R agonist that triggered a concentration-dependent transient increase in [Ca^2+^]_i_ in neutrophils (Fig 2A); the modulating effect was evident with propionate concentrations that were unable to induce an increase in [Ca^2+^]_i_ when added alone (Fig 2A). In contrast, no change in the Ca^2+^ response induced by P2Y_2_R agonist ATP was induced in neutrophils pre-treated with AZ1729 (Fig 2B). The response induced by propionate in AZ1729 sensitized neutrophils was inhibited by the FFA2R antagonist CATPB, but not by the P2Y_2_R antagonist AR-C118925 or the G*α*q selective inhibitor YM-254890 (Fig 2C). The two latter inhibited, however, the corresponding ATP induced response (Fig 2C). Taken together, these data show that AZ1729 selectively modulates the FFA2R-mediated transient rise in [Ca^2+^]_i_, and the sensitizing effect is the same as that recently described for the other allosteric FFA2R modulator Cmp58 (Lind, 2019a; Martensson et al., 2018).

**Figure 1.**
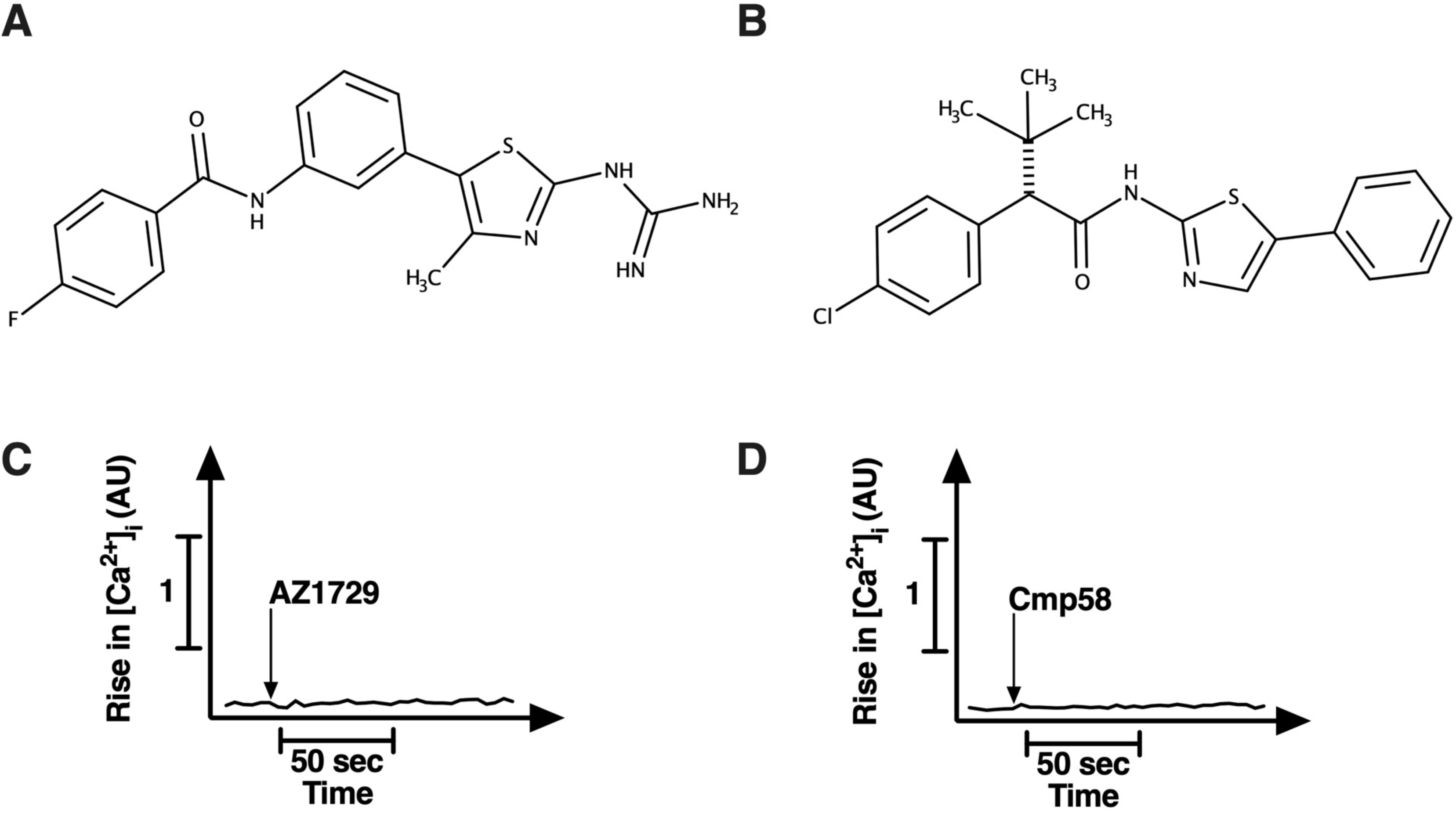
Structure of the FFA2R modulators AZ1729 and Cmp58 that lack the ability to trigger a rise in intracellular calcium in neutrophils. Chemical structure of two positive allosteric modulators for the free fatty acid receptor 2 (FFA2R); (**A**) AZ1729 and (**B**) Cmp58. Neutrophils loaded with Fura-2 were stimulated with AZ1729 (1 μM; **C**) or Cmp58 (1 μM; **D**) and the transient rise in intracellular Ca^2+^ [Ca^2+^]_i_ was measured continuously. The time point for addition of the respective modulator is indicated by an arrow. One representative experiment out of > 5 is shown. Abscissa, time of study (the bar represents 50 s); Ordinate, the increase in [Ca^2+^]_i_ is given as the ratio between Fura-2 fluorescence at 340 and 380 nm.

**Figure 2.**
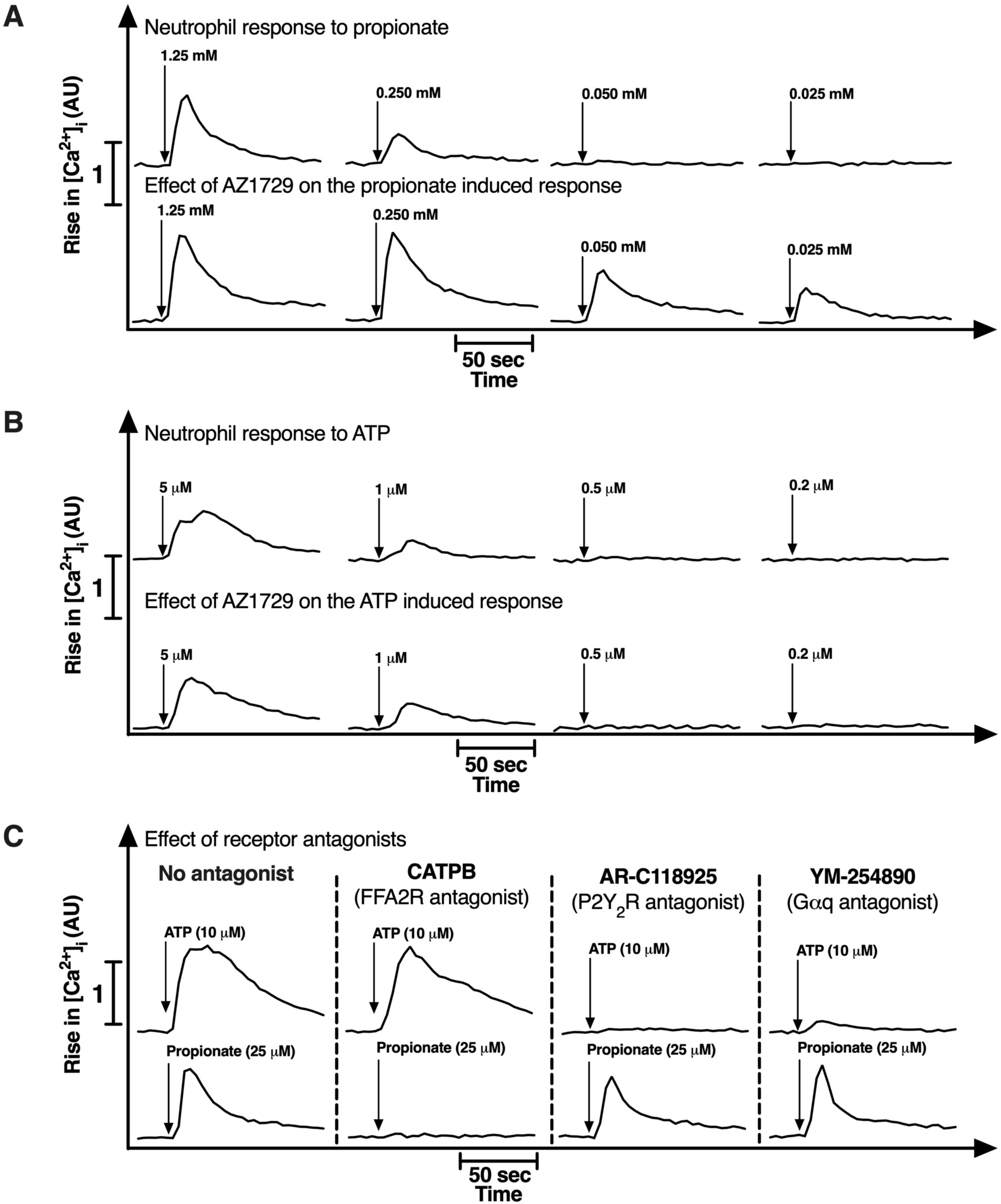
The allosteric FFA2R modulator AZ1729 selectively affects the transient rise in [Ca^2+^]_i_ in neutrophils when triggered by propionate. (**A**) Neutrophils were activated by different concentrations of propionate in the absence (upper panel) or presence of AZ1729 (incubated for 5 min with 1 μM AZ1729; lower panel), and the rise in [Ca^2+^]_i_ was determined. The time point for addition of propionate is indicated by arrows. (**B**) The same experimental setup was used with different concentrations of the P_2_Y2R agonist ATP as the activating agonist. The time point for addition of ATP is indicated by arrows. (**C**) Inhibition of the activation of AZ1729 sensitized neutrophil (5 min with 1 µM AZ1729) when triggered by ATP (10 μM, upper panel) or propionate (25 μM, lower panel). The time points for addition of ATP and propionate are indicated by arrows. The left parts of the panels show the responses without an inhibitor/antagonist, and then in order, the responses in the presence of the FFA2R antagonist, CATPB (100 nM), the P_2_Y2R antagonist AR-C118925 (1 μM) and, the Gαq inhibitor YM-254890 (200 nM), respectively. One representative experiment out of 3 is shown. Abscissa, time of study (the bar represents 50 s); Ordinate, the increase in [Ca^2+^]_i_ given as the ratio between Fura-2 fluorescence at 340 and 380 nm.

#### The allosteric modulator AZ1729 turns both propionate and ATP into highly efficient activators of the superoxide generating NADPH-oxidase

In agreement with the inability of AZ1729 to trigger a rise in [Ca^2+^]_i_, no activation of the superoxide generating NADPH-oxidase was induced by AZ1729 alone (Fig 3A), but in AZ1729 pre-incubated/sensitized cells, the non-activating orthosteric FFA2R agonist propionate (Lind, 2019a) was transferred to a potent activating agonist (Fig 3A). In quantitative terms the neutrophil superoxide production induced was not changed when the order of the ligands was reversed, i.e., when cells sensitized with propionate were activated by an addition of AZ1729 (not shown by figure). Although the P2Y_2_R agonist ATP triggered a Ca^2+^ response (Fig 2B and C), no neutrophil NADPH-oxidase activation was induced by ATP alone (Gabl et al., 2015; Lind, 2019a). However, when combined with AZ1729, ATP became a potent activator of the NADPH-oxidase (Fig 3A). In contrast to the response induced by two FFA2R ligands together (AZ1729 combined with propionate), the order in which AZ1729 and ATP was added could not be reversed; no superoxide release was induced by AZ1729 when ATP was added first (not shown by figure). To determine the concentration dependency of the propionate/ATP induced activation of neutrophils with their FFA2Rs allosterically modulated, we “clamped” the system by keeping the AZ1729 concentration constant (1 µM) and the priming (incubation) time with the modulator fixed to five minutes. Using this setup, it was clear that superoxide release from neutrophils activated with propionate was dependent on the concentration of the orthosteric FFA2R agonist with an EC_50_ value of ∼ 10 µM reaching a full response at ∼ 100 µM (Fig 3B). It was also clear that superoxide release from neutrophils activated with ATP was agonist concentration dependent with an EC_50_ value of <10 µM reaching a full response at ∼ 25 µM (Fig 3C). From a comparison of the sensitizing effects of AZ1729 and Cmp58, we could conclude that the latter was more potent (Fig 3D). In agreement with the earlier described allosteric modulation results obtained with Cmp58 (Lind, 2019a), the AZ1729/propionate response was inhibited by the FFA2R antagonist CATPB but not affected by the P2Y_2_R antagonist AR-C118925 or the G*α*q selective inhibitor YM-254890 (Fig 3E). The AZ1729/ATP response was inhibited by all three inhibitors/antagonists used (Fig 3E). These data show that the allosteric modulators affect not only binding and/or signaling triggered by orthosteric agonists, but affect also the sensitivity of FFA2R to the receptor cross-talk signals generated by ATP occupied P2Y_2_Rs.

**Figure 3.**
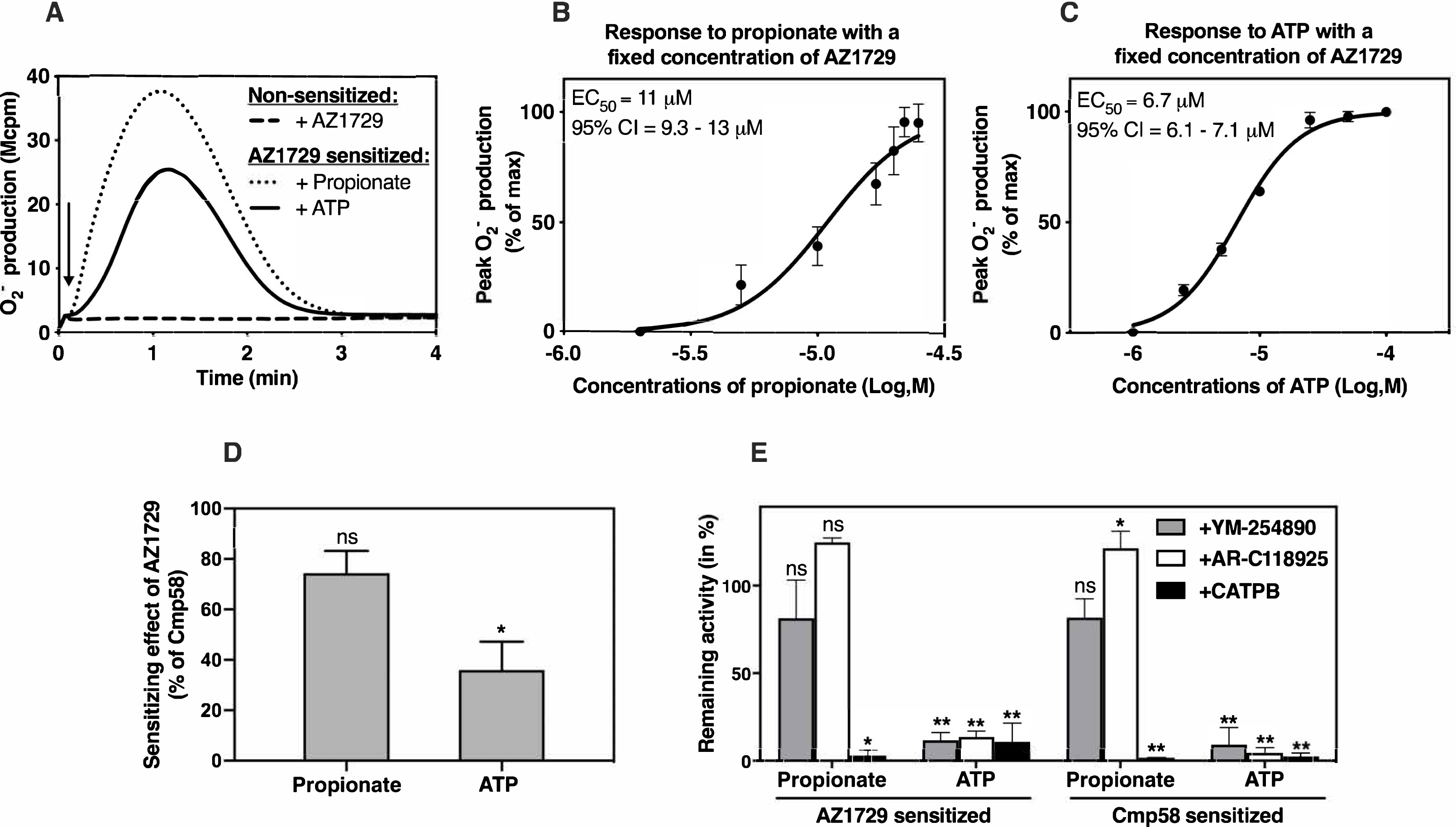
The allosteric FFA2R modulator AZ1729 affects the neutrophil NADPH-oxidase activity induced by propionate and ATP. (**A**) AZ1729 sensitized neutrophils (incubated for 5 min with 1 μM AZ1729) produce superoxide anions (O_2_^−^) generated by the NADPH-oxidase when activated with propionate (25 μM; dotted line) or ATP (10 μM; solid line). The production of O_2_^−^ was recorded continuously, and the time point for addition of the ligands is indicated by an arrow. The direct effect of AZ1729 (1 μM, dashed line) on non-sensitized neutrophils is shown for comparison, and one representative experiment out of > 5 is shown. (**B**) AZ1729 sensitized neutrophils (1 μM for 5 min) were activated by different concentrations of propionate as indicated and the peaks of O_2_^−^ production were determined. Abscissa, concentration of propionate; ordinate, O_2_^−^ production expressed in percent of the activity induced by 25 μM propionate (mean±SD, n = 3). (**C**) AZ1729 sensitized neutrophils (1 μM for 5 min) were activated by different concentrations of ATP as indicated and the peaks of O_2_^−^ production were determined. Abscissa, concentration of ATP; ordinate, O_2_^−^ production expressed in percent of the activity induced by 10 μM ATP (mean±SD, n = 3). (**D**) The peaks in O_2_^−^ production induced by propionate (25 μM) or ATP (10 μM) in AZ1729 sensitized neutrophils (1 μM for 5 min) were determined and compared to the responses induced by the same agonists in Cmp58 sensitized neutrophils (1 μM for 5 min). The sensitizing effect of AZ1729 is expressed as the ratio between the peak values (in percent) obtained with AZ1729 and Cmp58 as the sensitizing agent, respectively (mean±SD, n = 3). The statistical analysis was performed using paired Student’s t test comparing the peak responses from sensitized cells from AZ1729 or Cmp58 induced by propionate or ATP (ns=no significant difference). (**E**) Effects on O_2_^−^ production in AZ1729 and Cmp58 sensitized neutrophil (5 min with 1 µM of the respective modulator), of the FFA2R antagonist CATPB (100 nM), the P2Y2R antagonist AR-C118925 (1 μM) and the Gαq inhibitor YM-254890 (200 nM) when triggered by ATP (10 μM) or propionate (25 μM). The activity was determined as peak O_2_^−^ production values and the effects of the inhibitors expressed as remaining activity (in percent) of the response induced in Cmp58 and AZ1929 sensitized neutrophils in the absence of any inhibitor (mean±SD, n = 3). Statistical analyses were performed using a 1-way ANOVA comparing the peak responses in the absence and presence of respective inhibitor.

Taken together, we show that the sensitizing/modulating effects of AZ1729 on the NADPH-oxidase activity are qualitatively the same as that earlier described for the allosteric FFA2R modulator Cmp58 (Lind, 2019a), but quantitatively the activities induced using propionate and ATP concentrations giving a maximum response, were somewhat lower with AZ1729 than Cmp58 (Fig 3D). Although the effects mediated by AZ1729 and Cmp58 are very similar, they might possibly change the FFA2R signaling through interactions with different allosteric sites.

### When combined, the two non-activating allosteric FFA2R modulators AZ1729 and Cmp58 activate neutrophils and the signaling is functional selective

#### Activation of the superoxide (O_2_^−^)generating NADPH-oxidase

Allosteric modulators, that stabilize a receptor in a conformation that differs from that of the naive/resting receptor, possess a potential to change multiple receptor sites and by that affect a unique range of receptor triggered activities (Kenakin, 2017b). Accordingly, the structural change induced by an allosteric modulator could, in principle, affect not only the functional outcome mediated by an orthosteric agonist, but also that of an allosteric modulator that occupies another binding site on the same receptor. We, thus, set out to determine if there was an allosteric cooperativity between the two FFA2R modulators Cmp58 and AZ1729, and we found that AZ1729 sensitized (allosterically modulated) neutrophils were activated to produce O_2_^−^ not only when triggered by propionate (see Fig 3), but also when triggered by the other non-activating allosteric FFA2R modulator Cmp58 (Fig 4A). Together, the two allosteric FFA2R modulators AZ1729 and Cmp58 potently activated the NADPH-oxidase in neutrophils; the increased production/release of O_2_^−^ following activation was evident i) when the two modulators were added together (Fig 4A), ii) when Cmp 58 was used as the activating ligand and added to AZ1729 sensitized cells and, iii) when the order of the two was reversed and AZ1729 was used as the activating ligand and added to Cmp58 sensitized neutrophils (Fig 4B). The amount of O_2_^−^ produced by Cmp58/AZ1729 together, was comparable to that induced by propionate in the presence of one or the other of the allosteric modulators (Fig 4C). In quantitative terms the neutrophil superoxide production induced by 1µM Cmp58 in neutrophils pretreated for five minutes with 1µM AZ1729 was of the same magnitude as that induced by the orthosteric agonist propionate (Fig 4C). Further, the activity induced using a fixed concentration of Cmp58 as the sensitizing ligand was dependent on the concentration of AZ1729 with an EC_50_ value of around 100nM (Fig 4D). The activity induced using a fixed concentration of AZ1729 as the sensitizing ligand was dependent on the concentration of Cmp58 with an EC_50_ value of around 50nM (Fig 4E). In order to determine the allosteric effect by the combined activation by the two allosteric modulators, we “clamped” the activation system, using AZ1729 as the sensitizer (modulator) and Cmp58 as the activator. The concentration of the sensitizer was varied (1µM-0.01µM) and the priming (incubation) time was fixed to five minutes; using this setup, it was clear that superoxide release from neutrophils activated with Cmp58 was dependent on the concentration of the activator (Fig 5A) as well as the sensitizer (Fig 5B). The ligands produce a 2.2-fold decreased affinity for the receptor for each other and a 150- to 600-fold increased efficacy for each other when binding to their respective binding sites on the receptor. Both had a very low level of direct efficacy to induce response in the absence of the other modulator. The right hand part shows responses to Cmp58 in the absence and presence of a range of concentrations of AZ1729 (Fig 5B). It is evident that AZ1729 both increased the maximal response and produced sinistral shifts of the concentration-response curves to Cmp58 in a concentration dependent manner. Fitting of the complete set of curves to the equation used (see Material and Methods) yielded the allosteric parameters α = 0.45, β =150, K_B_ = 6.7 µM and τ_B_ = 0.2 Note that Cmp58 had a low level of direct efficacy, this was insufficient to produce visible response in the absence of AZ1729. The reciprocal analysis, namely responses in the presence of a range of concentrations of Cmp58 (Fig 5A) were also performed and the analysis yielded the allosteric parameters α = 0.4, β = 150, K_B_ = 6 μM, and τ_B_ = 0.2. The reciprocal sensitization mediated by the two allosteric modulators, represents a new regulatory mechanism in neutrophil that controls receptor signaling.

**Figure 4.**
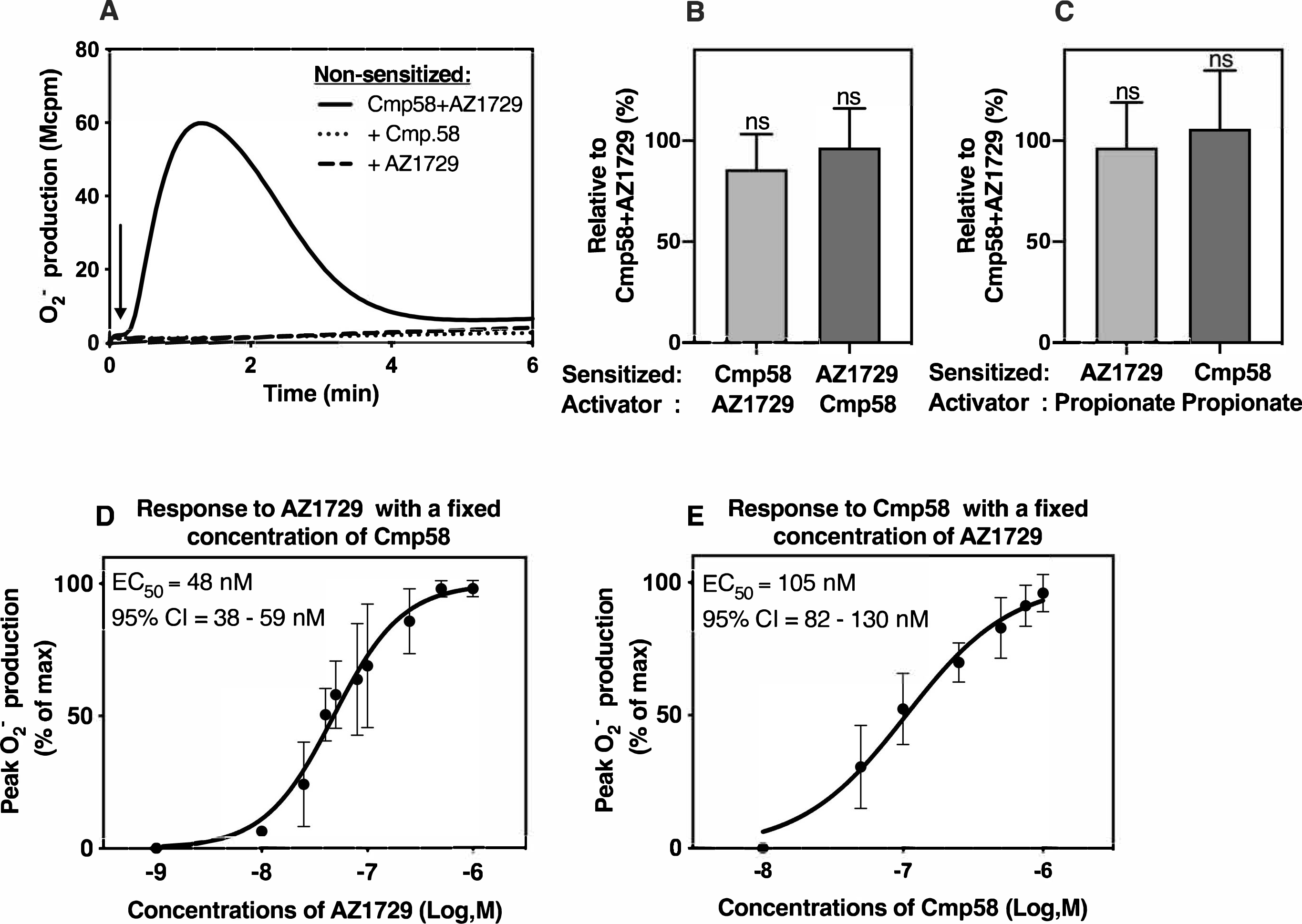
The two FFA2R modulators Cmp58 and AZ1729 together activate neutrophils to produce O_2_^−^. **(A)** Human neutrophils were stimulated with either of the two FFA2R modulators AZ1729 (1µM) and Cmp58 (1µM) that were added to neutrophils alone (dotted line for Cmp58 and dashed line for AZ1729) or together (1 μM each, solid line) and O_2_^−^ production was measured continuously. One representative experiment out of > 5 is shown and time for addition of the modulators is marked by an arrow. **(B)** Production of O_2_^−^ in AZ1729 and Cmp58 sensitized neutrophils (1µM for 5 min) when activated with the complementary allosteric modulator (1µM) compared to the response induced by the two allosteric modulators when added together (1 μM each) to non-sensitized neutrophils (peak responses in percent; mean±SD, n=4. Statistical analyses were performed using a 1-way ANOVA comparing the peak responses. **(C)** Production of O_2_^−^ in AZ1729 and Cmp58 sensitized neutrophils (1µM for 5 min) when activated with propionate (25µM) compared to the response induced by the two allosteric modulators when added together (1 μM each) to non-sensitized neutrophils (peak responses in percent; mean±SD, n=4). Statistical analyses were performed using a 1-way ANOVA comparing the peak responses when propionate was used as the activator and when the two allosteric modulators were added together. **(D**) Neutrophils sensitized with Cmp58 (1 μM for 5 min) were activated with different concentrations of AZ1729 as indicated. Superoxide production was recorded continuously, and the peak values of the responses were determined. Abscissa, concentration of propionate; ordinate, superoxide production expressed in percent of the activity induced by 1 μM AZ1729 (mean±SD, n = 3). (**E**) Neutrophils sensitized with AZ1729 (1 μM for 5 min) were activated with different concentrations of Cmp58 as indicated. Superoxide production was recorded continuously, and the peak values of the responses were determined. Abscissa, concentration of propionate; ordinate, superoxide production expressed in percent of the activity induced by 1 μM AZ1729 (mean±SD, n = 3).

**Figure 5.**
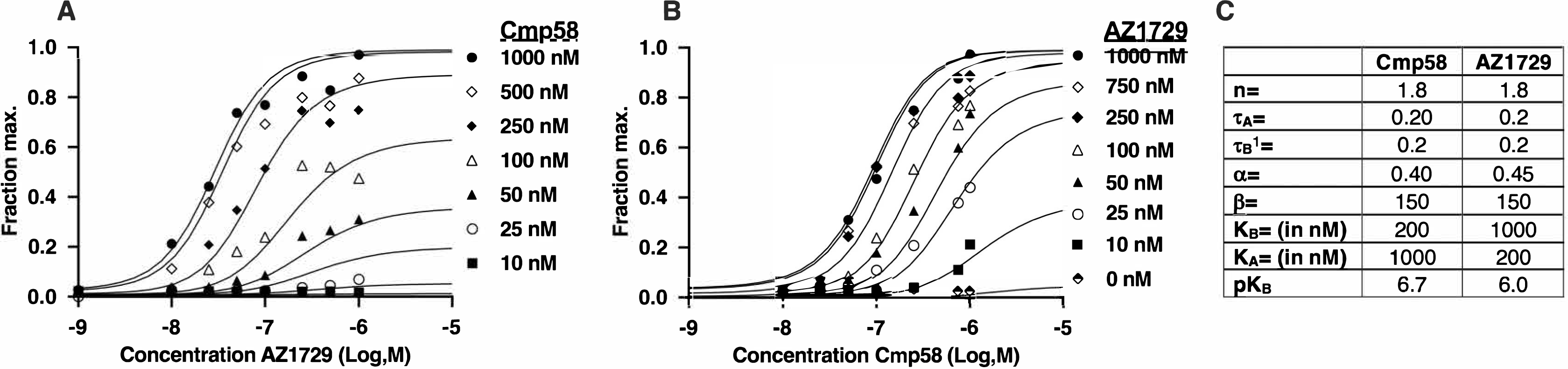
Analysis of allosteric effects using concentration response curves to AZ1729 and Cmp58. **(A)** Neutrophils were activated by difference concentrations of AZ1729 in presence of Cmp58 in concentrations of from 10 to 1000 nM. The neutrophil response was measured as superoxide production that was recorded continuously and the peak activities were used for the calculations. (**B)** Neutrophils were activated by difference concentrations of Cmp58 in presence of AZ1729 in concentrations up to 1000 nM. The neutrophil response was measured as superoxide production that was recorded continuously and the peak activities were used for the calculations. **(C)** Analysis of the data presented in the A and B parts of the figure (see above) yields values that characterize the allosteric interaction of [A] and [B] in a system independent manner. The allosteric effect is given by discrete values of α (effect on affinity), β (effect on efficacy), *K*_B_ (equilibrium dissociation constant of the modulator in the absence of A) and where applicable, τ_B_ (the efficacy of the modulator). What is required is the efficacy of the agonist (τ_A_), n (slope of the concentration-response curves) and some measure of the maximal response capability of the system (*E*_m_); these were obtained through fitting the concentration response curve to [A] in the absence of [B]. Fitting of the complete set of curves to the equation use yielded allosteric parameters for α, β, *K*_B_, and τ_B_. The values obtained in the reciprocal analysis, namely responses to Cmp58 in the presence of a range of concentrations of AZ1729, are also included. The values of τ_A_ and τ_B_ (supposedly the molecular efficacies of A and B, respectively) and the affinities of A and B for the receptor (given by the equilibrium dissociation constant K_A_ and K_B_) should be the same for each analysis. In addition, the values of α and β should also be the same since they simply describe the allosteric co-operativity between the two ligands A and B as they bind to the receptor. The experimental data was reasonably consistent with this constraint as seen by the relative values of α, β, K_B_, τ_A_, and τ_B_ used for the analyses. This suggests that a consistent allosteric effect is being quantified between Cmp58 and AZ1729.

#### The FFA2R selective antagonist CATPB inhibits neutrophil sensitization by allosteric FFA2R modulators

The FFA2R selective antagonist CATPB inhibits activation induced in non-sensitized neutrophils by conventional receptor agonists (Lind, 2019a; Martensson et al., 2018), suggesting that the antagonist blocks the othosteric binding site in FFA2R. As a consequence, it is not possible to use a measuring system that involves an orthosteric agonist as one of the ligands mediating the activation, in order to determine the direct inhibitory effects of the antagonist CATPB on the activity mediated by the allosteric FFA2R modulators. To determine the effects on the allosteric FFA2R sites that recognize AZ1729 and Cmp58, respectively, we used heterologous receptor sensitization as model system. Both modulators transfer ATP and low concentrations of the FPR1 agonist fMLF (Fig 6A and B) into NADPH-oxidase activating agonist, and it is evident from the results obtained with neutrophils sensitized with either Cmp58 or AZ1729 and activated with ATP, that the FFA2R antagonist CATPB inhibited the sensitization/modulation mediated separately by the allosteric modulators (Fig 3E). CATPB inhibited also the response induced by a concentration of fMLF, that when added to neutrophils in the absence of a modulator lacks activating effects (shown in the insets of Fig 6A with AZ1729 and in Fig 6B with Cmp58). For clarity, the fMLF induced response is inhibited not only by CATPB but also by cyclosporin H (an FPR1 antagonist; inset in Fig 6A and B). The FFA2R specific antagonist CATPB, thus blocks the modulating effects of Cmp58 and AZ1729, respectively, leading to an increased neutrophil response to ATP and fMLF. The FFA2R antagonist only partly inhibited the response induced by the two allosteric modulators together when using 1 µM concentrations of each, but the inhibitory effect was increased when the concentrations of the allosteric modulators were reduced (Fig 6C).

**Figure 6.**
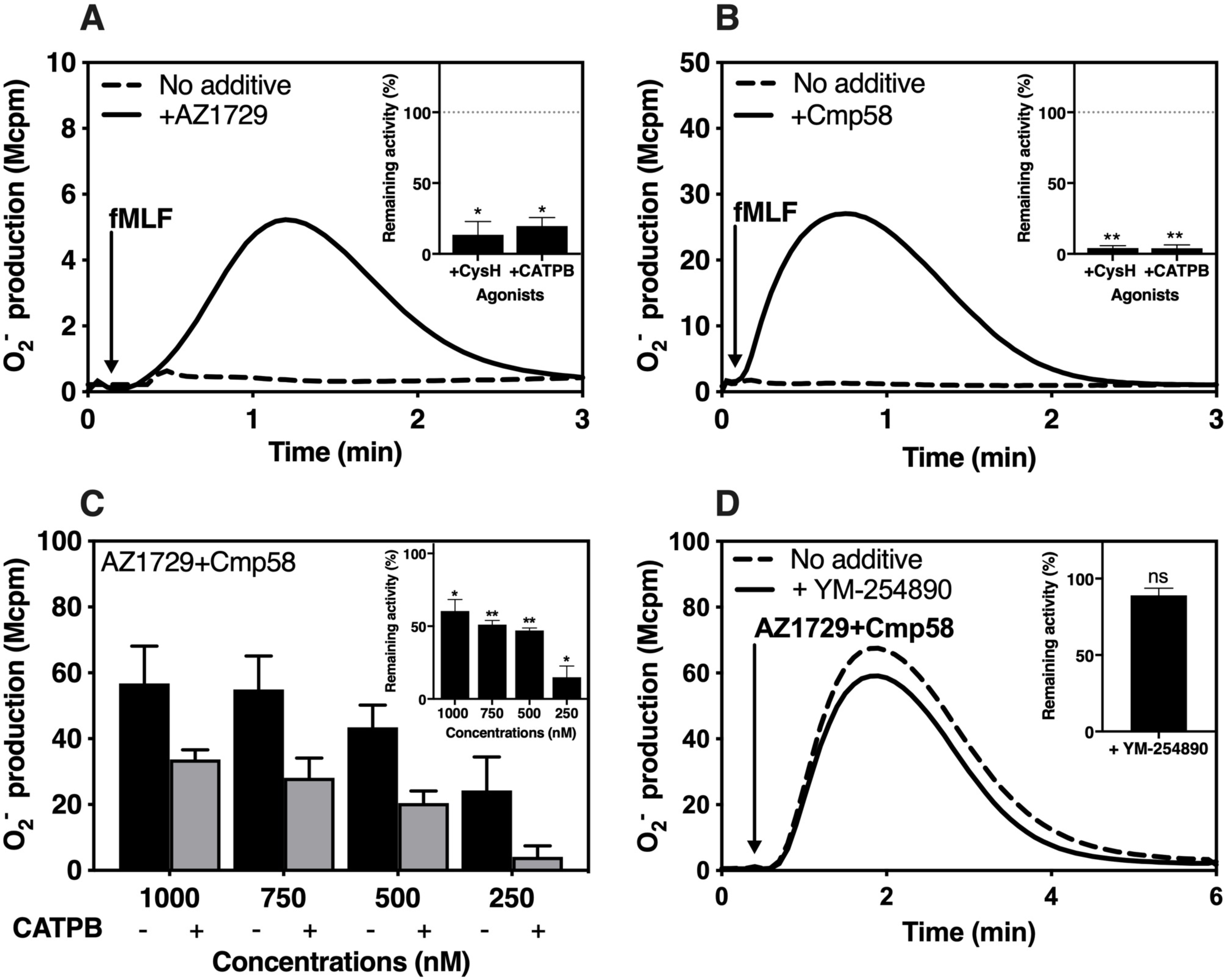
The FFA2R antagonist CATPB inhibits the effects of the FFA2R modulators. (**A**) Neutrophils incubated without (dashed line) and sensitized with AZ1729 (1µM for 5 min; solid line) were activated with the FPR1-specific peptide fMLF (1 nM) and the release of O_2_^−^ was measured. One representative experiment out of 3 is shown. Inset: The response induced by fMLF in AZ1729 sensitized (1µM for 5 min) neutrophils was inhibited by the FPR1 specific antagonist, CysH (1 μM) and the FFA2R specific antagonist CATPB (100 nM). The peak O_2_^−^ production values were determined and the ratios between the responses in the absence and presence of the respective antagonist were calculated and expressed in remaining activity (in percent) in the presence of the respective antagonist (mean±SD, n = 3). Statistical analyses were performed using a 1-way ANOVA comparing the peak responses in the absence and presence of respective inhibitor. (**B**) Neutrophils incubated without (dashed line) and sensitized with Cmp58 (1µM for 5 min; solid line) were activated with the FPR1-specific peptide fMLF (1 nM) and the release of O_2_^−^ was measured. One representative experiment out of 3 is shown. Inset: The response induced by fMLF in Cmp58 sensitized (1µM for 5 min) neutrophils was inhibited by the FPR1 specific antagonist, CysH (1 μM) and the FFA2R specific antagonist CATPB (100 nM). The peak O_2_^−^ production values were determined and the ratios between the responses in the absence and presence of the respective antagonist were calculated and expressed in remaining activity (in percent) in the presence of the respective antagonist (mean±SD, n = 3). Statistical analyses were performed using a 1-way ANOVA comparing the peak responses in the absence and presence of respective inhibitor. (**C**) Neutrophils incubated in the absence (black bars) or presence of the FFA2R antagonist CATPB (100 nM for 5 min; grey bars) were activated with equal but varying concentrations (as indicated) of Cmp58 and AZ1729 added together. Superoxide production was recorded continuously and the peak activities were determined and expressed in arbitrary units (mean±SD, *n*= 3). Inset: The inhibitory effects of CATPB on the neutrophil response induced by equal but varying concentrations of AZ1729+Cmp58 added together expressed as remaining activity induced by the respective AZ1729/Cmp58 concentration in the absence of the antagonist. The peak values were obtained from the main figure and the effects expresses as remaining activity in percent (mean±SD, n = 3). Statistical analysis was performed using paired Student’s t test comparing the peak responses obtained in the absence and presence, respectively, of the FFA2R antagonist. (**D**) Neutrophils incubated without (dashed line) or with the G*α*q specific inhibitor YM-254890 (200 nM for 5 min; solid line) were activated with AZ1729+Cmp58 (1 μM each) together, and the release of O_2_^−^ was recorded. One representative experiment out of 3 is shown. Time point for addition of AZ1729/Cmp58 is indicated by an arrow. Inset: The inhibitory effect of YM-254890 on the neutrophil response induced by AZ1729 and Cmp58 when added together, expressed as remaining activity (in percent of peak values) when compared with response induced of in the absence of the inhibitor (mean±SD, n = 3). The statistical analysis was performed using paired Student’s t test comparing the peak responses in the absence and presence of the inhibitor YM-254890.

It has been shown that allosterically modulated FFA2Rs have the capacity to signal through both G*α*i and G*α*q containing G-proteins (Bolognini et al., 2016), but the G*α*q selective inhibitor YM254890 was without effect on neutrophil activation when induced by AZ1729 and Cmp58 together, suggesting that G*α*q is not involved in signaling triggered by the two interdependent modulators (Fig 6D).

#### The cooperative neutrophil activation by the two interdependent allosteric FFA2R modulators occurs without any rise in [Ca^2+^]_i_

No direct neutrophil activating effects, measured as a change in [Ca^2+^]_i_, were induced by AZ1729 and Cmp58 alone, whereas the thresh-hold for the propionate response was lowered in the presence of either AZ1729 or Cpm58 (Fig 1 and 2; (Lind, 2019a)). It is clear from the data presented that when AZ1729 and Cmp58 were added together (or in sequence), they activate neutrophils to produce O_2_^−^, but this activation was achieved without any concomitant rise in [Ca^2+^]_i_ (Fig 7A). The inability to trigger a Ca^2+^ response was evident when the two modulators were added together, or added sequentially (Fig 7A).

**Figure 7.**
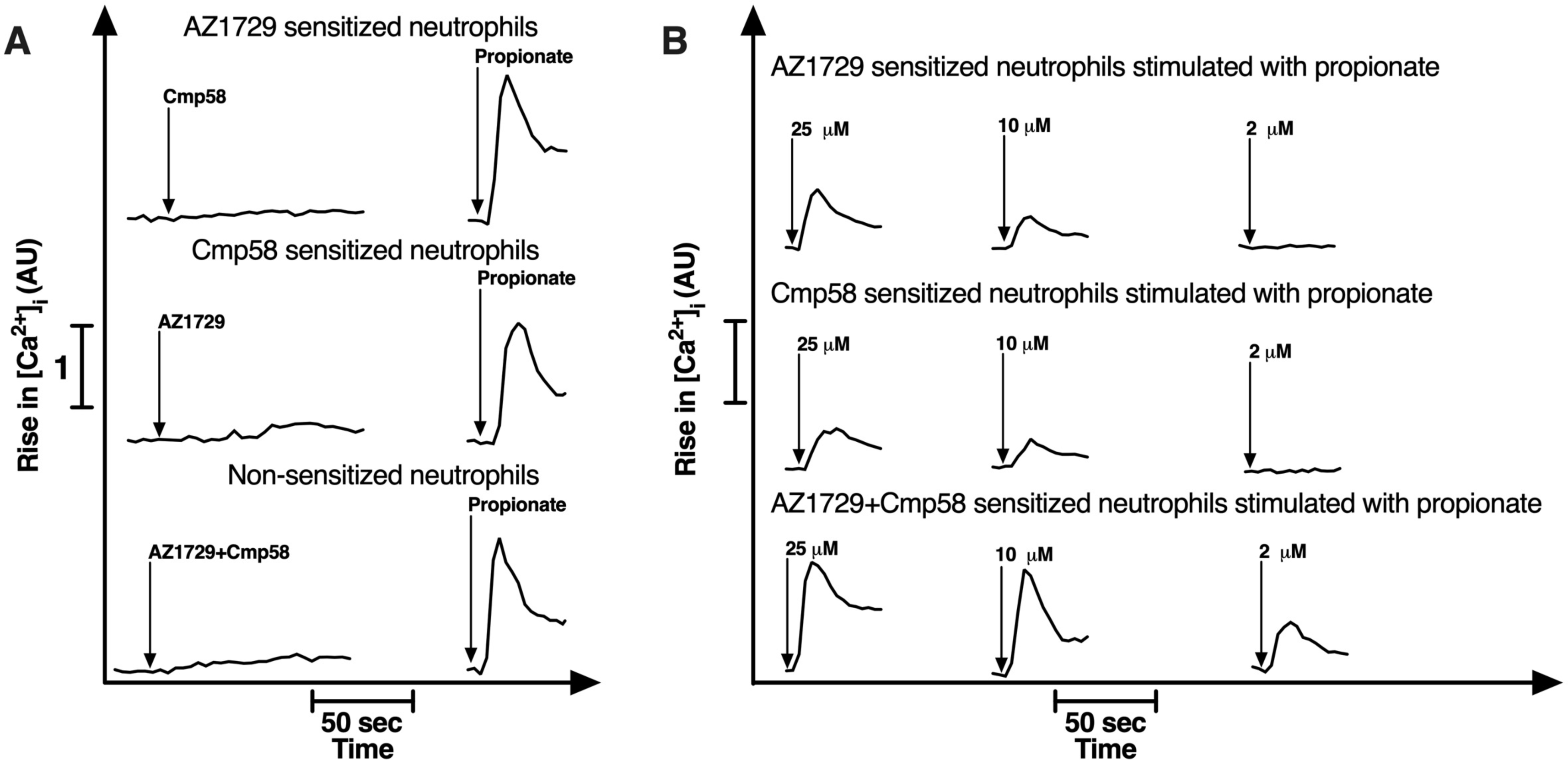
No transient rise in [Ca^2+^]_i_ is triggered when the allosteric modulators are combined and the sensitizing effect on the propionate response is augmented. (**A**) Neutrophils sensitized with AZ1729 (1 μM 5 min; upper left panel) and Cmp58 (1µM 5 min; middle left panel) were activated with Cmp58 (1 μM, upper left panel) and AZ1729 (1µM, middle left panel). Non-sensitized neutrophils (lower left panel) were activated with AZ1729+Cmp58 (1 μM each) added together. Propionate (25µM; right panel) was added to the cells shown in the left panel 2 min after the time point (indicated by arrows) for addition of the respective allosteric modulator. One representative experiment out of 3 independent experiment is shown. Abscissa, time of study (the bar represents 50 s); Ordinate, the increase in [Ca^2+^]_i_ is expressed as the ratio between Fura-2 fluorescence at 340 and 380 nm. (**B**) Neutrophils sensitized (2 min) with AZ1729 (1µM, upper panel) Cmp58 (1µM, middle panel) and AZ1729/Cmp58 together (1µM each, lower panel) were subsequently activated by different concentrations of propionate (as indicated). One representative experiment out of 3 independent experiment is shown. Abscissa, time of study (the bar represents 50 s); Ordinate, the increase in [Ca^2+^]_i_ expressed as the ratio between Fura-2 fluorescence at 340 and 380 nm.

#### Biased desensitization in neutrophils mediated by cooperative action of the two allosteric FFA2R modulators AZ1729 and Cmp58

The NADPH-oxidase activity induced by the cooperative action of AZ1729 and Cmp58 was very pronounced with a peak of activity reached in around a minute and the response was then the rapidly terminated; these neutrophils were homologously desensitized as no response was induced when a new dose of AZ1729/Cmp58 was added (Fig 8 A, Inset). Propionate induced, however, a response in AZ1729/Cmp58 sensitized neutrophils, but the response was small compared to that induced by propionate in solely AZ1729 or Cmp58 sensitized neutrophils (Fig 8A and B). The response induced by the FPR1 agonist fMLF (100nM) was not affected by the allosteric FFA2R modulators alone or in combination (Fig 8C and D). Neutrophils sensitized with AZ1729/Cmp58 together (or in sequence) retained the ability to respond with a rise in [Ca^2+^]_i_ when triggered by propionate (Fig 7A). In fact, a response was induced in these cells even with propionate concentrations unable to trigger a response in neutrophils sensitized with either of the allosteric FFA2R modulators alone (Fig 7B). Taken together these data suggest that the Cmp58/AZ1729-induced desensitization is functional selective, favoring the Ca^2+^over the NADPH-oxidase activating pathway.

**Figure 8.**
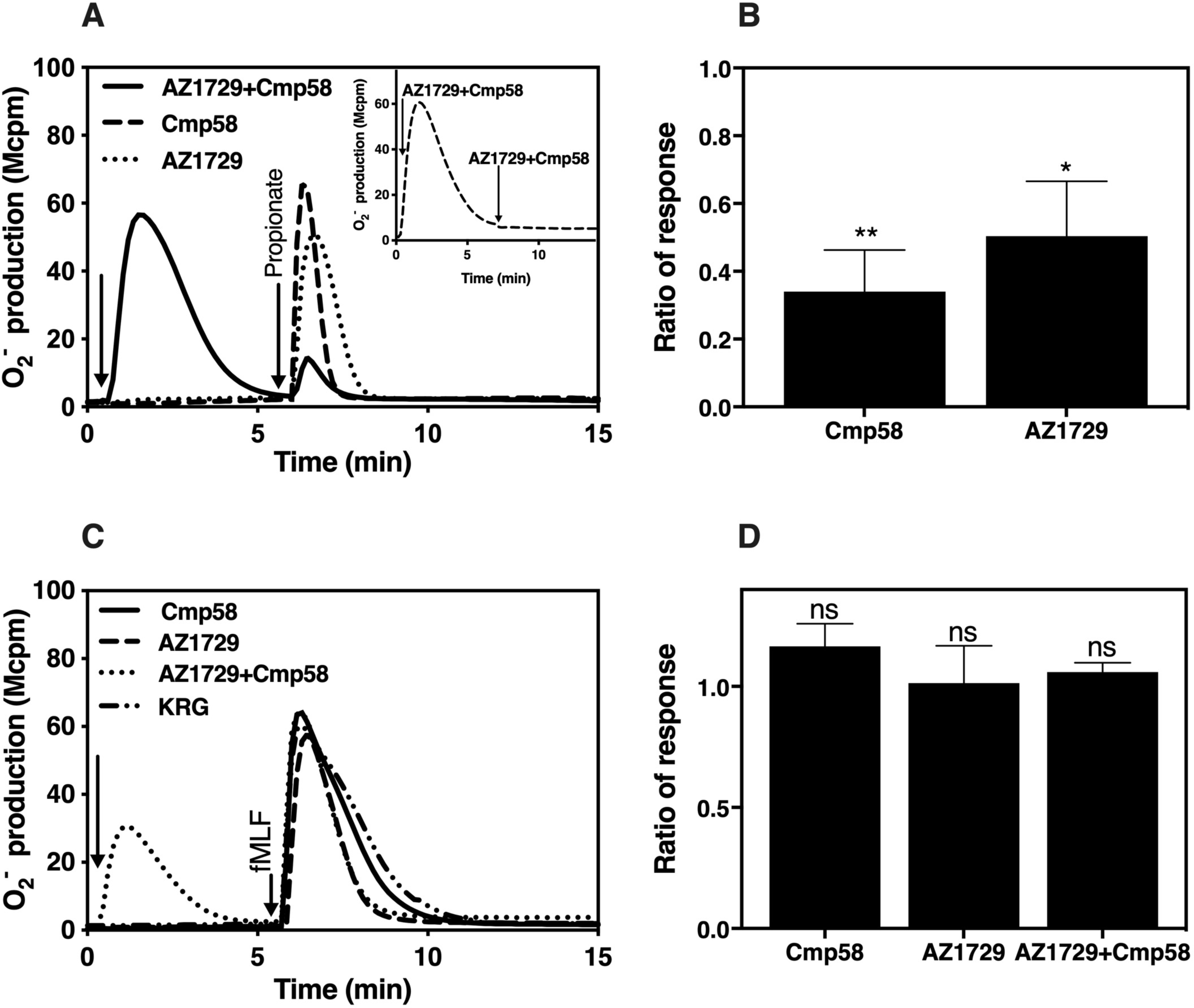
The propionate induced production of O_2_^−^ is reduced in neutrophils activated by AZ1729 and Cmp58 together. (**A**) Neutrophils were sensitized with AZ1729 (1µM, dotted line) or Cmp58 (1µM, dashed line) or activated by the two together (1µM each, solid line); the ligands were added at the time point indicated by the arrow to the left, and at the time point indicated by the arrow to the right, propionate (25µM) was added to the cells, and the production of O_2_^−^ was determined continuously. Inset: Neutrophils activated by Cmp58+AZ1729 together (1 μM each, added at the arrow to the left) were once again activated with the same ligands (1µM each, added at the time point indicated by the arrow to the right). One representative of 5 independent experiments is shown. (**B**) The NADPH-oxidase activity induced by propionate (25 μM) in AZ1729+Cmp58 (1 μM each) activated neutrophils, compared to the response induced by propionate in neutrophils sensitized with the allosteric modulators alone, calculated from the peak values of responses induced in AZ1929 and Cmp58 sensitized and AZ1729/Cmp58 activated neutrophils and expressed as the ratios between the responses in activated (with AZ1729+Cmp58) and sensitized neutrophils (with AZ1729 or Cmp58), respectively (mean±SD, n=5). Statistical analyses were performed using a 1-way ANOVA comparing the peak responses obtained when propionate was used to activate neutrophils sensitized with either of the two allosteric modulators alone and together. (**C**) The same experimental set up as that described in (**A**) was used but propionate was replaced by fMLF (100nM). One representative experiment out of 3 is shown. (**D**) The NADPH-oxidase activity induced by fMLF (100 nM) in AZ1729/Cmp58 (1 μM each) activated neutrophils and in neutrophils sensitized with the allosteric modulators alone, calculated from the peak values of responses induced in AZ1929 and Cmp58 sensitized and AZ1729/Cmp58 activated neutrophils and expressed as the ratios between the response in non-sensitized/activated neutrophils and activated (with AZ1729/Cmp58) or sensitized neutrophils (with AZ1729 or Cmp58), respectively (mean±SD, n=3). Statistical analyses were performed using a 1-way ANOVA comparing the peak responses obtained when fMLF was used to activate neutrophils sensitized with either of the two allosteric modulators alone and together.

## Discussion

The allosteric modulator AZ1729 primes neutrophils in their response to the orthosteric FFA2R agonist propionate and heterologously sensitize neutrophils to P2Y_2_R and FPR agonists, results in line with earlier findings obtained with another allosteric FFA2R modulator, Cmp58 (Lind, 2019a). The experimental data presented also support the earlier described receptor-cross-talk signaling mechanism as the molecular bases for priming of the ATP and fMLF induced activation of the neutrophil NADPH-oxidase (see model in Fig 9A; (Lind, 2019b; Onnheim et al., 2014)). One of the new findings presented, is that the modulator AZ1729 turns the other non-activating allosteric modulator Cmp58 into a potent neutrophil activating molecule and, as required by allosteric theory, this activation is reciprocal, that is Cmp58 also turns AZ1729 into a potent neutrophil activating ligand. Another novel finding is that the down-stream signaling profile of FFA2R, when activated by the two allosteric modulators, is biased in that no transient rise in intracellular Ca^2+^ is triggered during activation by the two allosteric modulators together, and the desensitized state is functional selective in that propionate induced activation of the superoxide generating NADPH oxidase is reduced whereas the PLC-PIP_2_-IP_3_-Ca^2+^ signaling pathway is potentiated (see the suggested model in Fig 9B and C).

**Figure 9.**
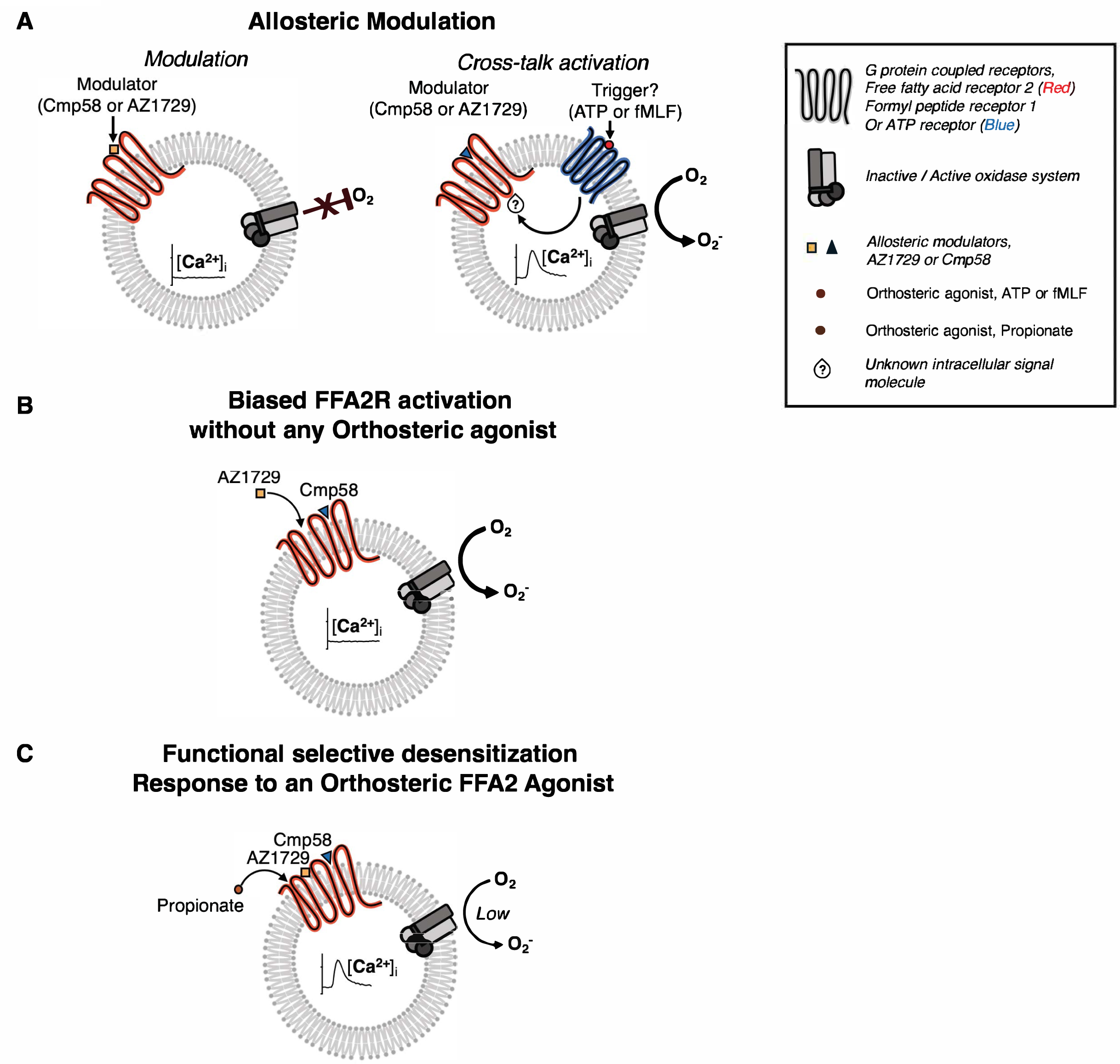
Model for the allosteric modulation as the basis for receptor cross talk, biased FFA2R activation without any orthosteric agonist and functional selective FFA2R desensitization. **A. Allosteric FFA2R modulation**: Left: No signals are generated by FFA2R expressed in the neutrophil plasma membrane by either of the allosteric FFA2R modulators Cmp58 or AZ1729 in the absence of a ligand. **Receptor cross talk activation:** Right: Binding of either of the modulators Cmp58 and AZ1729, opens for a receptor cross-talk between P_2_Y2R/FPR1 and FFa2R. The signals that trigger a transient rise in [Ca^2+^]_i_ and an activation of the ROS producing NADPH-oxidase starts down-stream of the agonist occupied P_2_Y2R or FPR1 and G-protein dependent signals generated down-stream of P2Y_2_R/FPR1 activate the allosterically modulated FFA2R. **B. Biased FFA2R activation without any orthosteric agonist.** The allosteric FFA2R modulator AZ1729 transfers another allosteric modulator,Cmp58, into a potent activator of the neutrophil ROS-producing NADPH-oxidase. The effect of the two modulators is reciprocal and functional selective, in that the oxidase is activated without any rise in the concentration of [Ca^2+^]_i_. **C. Functional selective desensitization of FFA2R.** Neutrophils synergistically activated by and homologously desensitized to the two interdependent allosteric FFA2R modulators AZ1729 and Cmp58, can be activated by the orthosteric FFA2R agonist propionate, but the response is functional selective; whereas the ROS producing capacity by propionate is reduced, the ability to trigger a transient rise in [Ca^2+^]_i_ is potentiated.

AZ1729 has been shown to act both as an FFA2R agonist and as a positive FFA2R modulator but this compound as well Cmp58, identified initially in screening studies designed to identify FFA2R ligands (Bolognini et al., 2016; Wang et al., 2010), lack the carboxylic acid entity (Fig 1) suggested to be needed for the activities induced by orthosteric FFA2R agonists (Sergeev et al., 2016). This implies that they both are ligands that interact with allosteric receptor sites, which is in agreement with the fact that AZ1729 and Cmp58 alone have no detectable direct neutrophil activating effects and the two exert very similar modulating functions with respect to their effects on the response induced by the othosteric FFA2R agonist propionate and also on the receptor cross-talk with the agonist occupied P2Y_2_Rs/FPRs. The precise receptor sites involved in the modulation are not known, but the fact that one modulator turns the other into a potent neutrophil activating agonist, suggests that the two interact with different (but possibly overlapping) allosteric binding sites. The orthosteric FFA2R antagonist CATPB (Milligan et al., 2017) inhibits not only the activity induced by the orthosteric agonist propionate, but also the response induced in sensitized neutrophils activated by ATP and low concentrations of fMLF, an activation that totally depends on the presence of either of the allosteric FFA2R modulators. The modulating effects of both AZ1729 and Cmp58 are, thus, achieved through their interaction with FFA2R and the inability of the two allosteric modulators to activate the PLC-PIP_2_-IP_3_-Ca^2+^ signaling rout, suggests that their respective FFA2R binding site is distinct from that of the orthosteric agonist propionate, but also that all three binding sites are close enough to be inhibited by the orthosteric FFA2R antagonist CATPB.

The signals generated by the orthosteric agonist propionate in neutrophils sensitized by the allosteric FFA2Rs modulated by AZ1729 or Cmp58, are balanced in the sense that the activation of the superoxide anion producing NADPH-oxidase is accompanied by a transient rise in [Ca^2+^]_i_. This signaling (being insensitive to the Gαq inhibitor) is possibly mediated through Gαi, but to verify this, experiments with a Gαi selective inhibitor are required, yet no such inhibitor is available. Pertussis toxin is assumed to be a Gαi selective inhibitor, but in neutrophils this toxin inhibits signaling also by receptors that are Gαq-coupled (Bylund et al., 2003; Dahlgren, 1987; Forsman et al., 2013; Gabl et al., 2015; Holdfeldt et al., 2017). The signals generated by FFA2R when triggered by AZ1729 and Cmp58 together, induce an assembly and activation of the neutrophil NADPH-oxidase without any transient rise in [Ca^2+^]_i_. These data support earlier findings that there is no direct link between the signals that activate the neutrophil NADPH-oxidase and the Ca^2+^ signaling route (Bylund et al., 2003; Dahlgren, 1987; Forsman et al., 2013; Gabl et al., 2015), but more importantly, the conformational change induced during activation of allosterically modulated FFA2Rs differs when the activating ligand occupies the orthosteric binding site and an allosteric site, respectively. The precise signals that leads to an assembly and activation of the superoxide generating oxidase is not known, but based on the facts that irrespectively if the response is balanced (as with propionate) or functional selective (as with an allosteric modulator), no inhibition was obtained with the G*α*q inhibitor YM254890. Hence, we suggest that the signals leading to an activation of the oxidase are generated independent of the G-protein-PLC PIP_2_-IP_3_-signaling pathway.

It should also be noticed, that following the termination of the response induced by AZ1729 and Cmp58 together, the receptors are non-responsive (homologously desensitized) to a new dose of these ligands, but still responsive to the orthosteric FFA2R agonist propionate. Moreover, the response induced by propionate in neutrophils first activated by the two allosteric modulators, was functionally selective with a pronounced/amplified Ca^2+^ response but a very weak superoxide production, suggesting a novel receptor biased desensitization mechanism. Currently, very little is really known about GPCR desensitization mechanisms in neutrophils, but in these cells, a physical coupling of the desensitized receptor to the actin cytoskeleton, rather than to β-arrestin, has been suggested to play an important role in homologous receptor desensitization (Dahlgren et al., 2016). The basis for this has been proposed to involve a physical separation of the receptor from the signaling G-protein achieved through a coupling of the agonist occupied receptors to the actin cytoskeleton(Jesaitis and Klotz, 1993), but there are also additional desensitization mechanisms that remains to be characterized in detail (Gabl et al., 2015). Irrespective of the precise modulation mechanism for the allosteric modulators (that is lowered energy-barrier for the conformational shift to a signaling state or a higher affinity for the second ligand), the outcome of the modulating effect of AZ1729 in neutrophils is that Cmp58 is turned into a potent biased activating ligand that triggers neutrophils to generate and release superoxide anions, and the activation achieved is consistent with an interdependent allosteric synergistic function of AZ1729 and Cmp58. It should also be noticed, that the effect on the induced response was the same as when the order of addition was reversed and Cmp58 was used as sensitizing ligand. Allosteric interactions should be reciprocal in nature and the results presented are thus in agreement with the reciprocity characterizing the effects of allosteric modulators (Kenakin, 2017a; Kenakin, 2017b). The basic characterization of AZ1729 clearly show that this FFA2R regulatory compound interacts with an allosteric site in the receptor and it has the capacity to act both as activating allosteric agonist with a signaling insensitive to G*α*q inhibition, and as a positive allosteric modulator that primes the receptors/cells when activated by an orthosteric agonist (Bolognini et al., 2016). The signaling that activate allosterically modulated FFA2Rs can at this point only be speculative, but the concept of biased signaling/functional selectivity is now firmly established as an important mechanism that regulates GPCR mediated activities also in neutrophils. A major limitation of the possibilities to elucidate the relevance thereof in these cells has been the lack of characterized receptor-specific agonists with different signaling properties. Our data generated from this study clearly demonstrate that the biased signaling/functional selectivity concept is valid for FFA2R, a neutrophil-expressed receptor important for the regulation of inflammatory processes (Gabl et al., 2015), and the data presented provide direct evidence for biased FFA2R signaling, resulting in an activation of the neutrophil NADPH-oxidase without involvement of the PLC-PIP_2_-IP_3_-Ca^2+^-route. In addition, a novel functionally selective desensitization mechanism is shown, that allows the orthosteric agonist to initiate one signaling pathway but not another – the desensitized state being functional selective in that propionate induced activation of the superoxide generating NADPH oxidase is reduced whereas the PLC-PIP_2_-IP_3_-Ca^2+^ signaling pathway is potentiated.

## Authorship contributions

*Participation in research design*: Lind, Holdfeldt, Mårtensson, Sundqvist, Björkman

*Conducted experiments:* Lind, Holdfeldt, Sundqvist

*Contributed analytical tools:* Kenakin

*Performed data analysis:* Lind, Holdfeldt, Kenakin, Forsman, Dahlgren

*Planned and supervised the research and wrote the manuscript:* Forsman, Dahlgren

*Contributed to the writing of the manuscript:* Lind, Holdfeldt, Martensson, Sundqvist, Bjorkman, Kenakin

## Footnotes

The work was supported by the Swedish Medical Research Council (CD,005601; HF, 02448) the King Gustaf V 80-Year Foundation (CD FAI 2014-0011), the Clas Groschinsky Foundation (HF, MI562), the Wilhelm and Martina Lundgrens Scientific Foundation (MS 2019-3102), Magnus Berwall Foundation (MS, 2018-02579), the Swedish Foundation for Strategic Research (HF, SM-17-0046, Åke Wibergs Foundation (HF, M15-005), and the Swedish state under the ALF-agreement (CD, ALFGBG 72510).

We thank Karolina Nilsson at AstraZeneca for kindly providing the allosteric FFA2R modulator AZ1729 and we also thank the members of the Phagocyte Research Group at the Sahlgrenska Academy, University of Gothenburg, for critically discussing the results and the manuscript. The experimental work performed by Linda Bergqvist is also gratefully acknowledged

